# A novel class of long small RNAs associates with Argonaute1 and is up-regulated by nutrient deprivation in the alga *Chlamydomonas*

**DOI:** 10.1101/2022.03.17.484771

**Authors:** Yingshan Li, Eun-Jeong Kim, Adam Voshall, Etsuko N. Moriyama, Heriberto Cerutti

## Abstract

Small RNAs (sRNAs) associate with Argonaute (AGO) proteins forming effector complexes with key roles in gene regulation and defense responses against molecular parasites. In multicellular eukaryotes, extensive duplication and diversification of RNA interference (RNAi) components have resulted in intricate pathways for epigenetic control of gene expression. The unicellular alga *Chlamydomonas reinhardtii* also has a complex RNAi machinery, including three AGOs and three Dicer-like (DCL) proteins. However, little is known about the biogenesis and function of most endogenous sRNAs. We demonstrate here that *Chlamydomonas* contains uncommonly long sRNAs (>26 nt), which associate preferentially with AGO1. Somewhat reminiscent of animal PIWI-interacting RNAs, these long sRNAs are derived from moderately repetitive genomic clusters and their biogenesis appears to be Dicer-independent. Interestingly, long sRNA encoding sequences have been conserved and amplified in phylogenetically related *Chlamydomonas* species. Additionally, expression of several long sRNAs increases substantially under nutrient deprivation, correlating with the downregulation of predicted target transcripts. We hypothesize that the transposon-like sequences encoding long sRNAs might have been ancestrally targeted for silencing by the RNAi machinery but, during evolution, some long sRNAs might have fortuitously acquired endogenous target genes and become integrated into gene regulatory networks.

## Introduction

Small silencing RNAs (sRNAs) play important roles in, among other processes, regulation of gene expression, heterochromatin formation, DNA methylation, maintenance of genome stability, intercellular communication, transposon repression and/or defense against viruses in a wide range of eukaryotes (Ghildiyal and Zamore, 2009; Borges and Martienssen, 2015, Wendte and Pikaard, 2017; Yu et al., 2017; Bartel, 2018; Ozata el al., 2019; Chen and Rechavi, 2021). Endogenous sRNAs are generally 20-30 nucleotides (nts) in length and associate with members of the Argonaute (AGO) family of proteins, forming effector complexes (Ghildiyal and Zamore, 2009; Swarts et al., 2014; Borges and Martienssen, 2015, Wendte and Pikaard, 2017; Yu et al., 2017; Bartel, 2018; Ozata el al., 2019; Chen and Rechavi, 2021; Iwakawa and Tomari, 2022). These effector complexes then recognize target sequences, by complementarity to the guide sRNAs, and trigger post-transcriptional or transcriptional gene silencing by diverse molecular mechanisms. Beyond these defining features, many different classes of sRNAs have been described in a broad spectrum of eukaryotes (Katiyar-Agarwal et al., 2007; Ghildiyal and Zamore, 2009; Axtell, 2013; Borges and Martienssen, 2015; Yu et al., 2017; Bartel, 2018; Hardcastle et al., 2018; Ozata el al., 2019; Feng et al., 2020; Lunardon et al., 2020; Müller et al., 2020; Rzeszutek and Betlej, 2020; Wu et al., 2020; Alves and Nogueira, 2021; Chen et al., 2021; Chen and Rechavi, 2021).

In most species, endogenous sRNAs derived from double-stranded RNA (dsRNA) precursors processed by Dicer-like (DCL) endonucleases can be grouped into two main classes: microRNAs (miRNAs) and small interfering RNAs (siRNAs) (Ghildiyal and Zamore, 2009; Axtell, 2013; Borges and Martienssen, 2015; Yu et al., 2017; Bartel, 2018; Lunardon et al., 2020; Müller et al., 2020; Chen et al., 2021). miRNAs are typically 21-22 nt long, processed from imperfectly paired stem-loop regions of single-stranded RNA (ssRNA) precursors, and regulate gene expression through mRNA degradation and/or translation repression (Ghildiyal and Zamore, 2009; Axtell, 2013; Borges and Martienssen, 2015; Yu et al., 2017; Bartel, 2018). siRNAs are produced from near-perfectly complementary dsRNAs of various origins, participate in post-transcriptional or transcriptional gene silencing, and are grouped in multiple subclasses (Ghildiyal and Zamore, 2009; Axtell, 2013; Borges and Martienssen, 2015; Lunardon et al., 2020; Chen et al., 2021; Chen and Rechavi, 2021). In angiosperms, the most abundant siRNA subclass, ∼24 nt heterochromatic siRNAs, is involved in the canonical RNA-directed DNA methylation pathway primarily targeting transposons and other repeats (Axtell, 2013; Borges and Martienssen, 2015; Lunardon et al., 2020; Chen et al., 2021; Chen and Rechavi, 2021). Another major siRNA subclass consists of secondary siRNAs, including trans-acting siRNAs, phased siRNAs and epigenetically activated siRNAs, which may silence genes and/or transposons. Their biogenesis is generally triggered by miRNA-directed cleavage of a transcript, which is then converted to dsRNA by an RNA-dependent RNA polymerase and processed by DCL proteins into siRNAs, often in a phased pattern (Axtell, 2013; Borges and Martienssen, 2015; Lunardon et al., 2020; Chen et al., 2021; Chen and Rechavi, 2021). Several additional siRNA subtypes (e.g., natural antisense transcript-derived siRNAs, hairpin-derived siRNAs, DNA double-strand break-induced small RNAs, and bacteria-induced long siRNAs) have also been described in land plants, particularly in model species such as *Arabidopsis thaliana*, and the boundaries among the siRNA classes are becoming quite diffuse (Katiyar-Agarwal et al., 2007; Axtell, 2013; Borges and Martienssen, 2015; Hardcastle et al., 2018; Lunardon et al., 2020; Rzeszutek and Betlej, 2020; Wu et al., 2020; Alves and Nogueira, 2021; Chen and Rechavi, 2021). Metazoans have an additional large class of longer sRNAs (∼22-35 nt) termed PIWI-interacting RNAs (piRNAs). piRNAs are distinct from siRNAs since they derive from ssRNAs, do not require DCL proteins for their processing and bind to PIWI (P-element Induced Wimpy Testis) proteins, an evolutionarily distinct clade of Argonaute proteins (Ghildiyal and Zamore, 2009; Swarts et al., 2014; Ozata el al., 2019).

It is generally accepted that sRNA-mediated silencing evolved as a defense mechanism against viruses and transposable elements and was later adapted to regulate the expression of endogenous genes (Cerutti and Casas-Mollano, 2006; Shabalina and Koonin, 2008). Extensive duplication and specialization of proteins involved in sRNA biogenesis and/or effector functions have contributed to pathway diversification (Ghildiyal and Zamore, 2009; Swarts et al., 2014; Borges and Martienssen, 2015; Lee and Carroll, 2018; Wang et al., 2021; Chen and Rechavi, 2021); and the sorting of sRNAs into specific AGOs/effector complexes ultimately determines their biological function(s) (Ghildiyal and Zamore, 2009; Czech and Hannon, 2011; Borges and Martienssen, 2015; Iwakawa and Tomari, 2022). In land plants, the structure of sRNA duplex precursors (such as thermodynamic asymmetry), the presence and location of mismatches and bulges, as well as the identity of the 5’-terminal nucleotide of sRNAs affect their loading onto individual AGOs and the choice of guide strand (Mi et al., 2008; Takeda et al., 2008; Havecker et al., 2010; Czech and Hannon, 2011; Zhu et al., 2011; Frank et al., 2012; Endo et al., 2013; Zhang et al., 2014; Borges and Martienssen, 2015; Iwakawa and Tomari, 2022). For instance, *A. thaliana* has 10 Argonautes and AGO1 and AGO10 preferentially bind sRNAs with a 5’-uracil, AGO5 has a bias for 5’-cytosine, whereas AGO2, AGO4, AGO6, and AGO9 prefer 5’-adenine (Mi et al., 2008; Takeda et al., 2008; Havecker et al., 2010; Czech and Hannon, 2011; Frank et al., 2012; Borges and Martienssen, 2015). The resulting large variety of sRNA-directed effector complexes function in distinct, yet often intertwined, regulatory pathways that influence development, responses to both abiotic and biotic stresses, reproduction, and genome reprogramming (Ghildiyal and Zamore, 2009; Czech and Hannon, 2011; Borges and Martienssen, 2015; Lee and Carroll, 2018; Chen and Rechavi, 2021). Moreover, new sRNA classes and new members of existing classes continue to be discovered, sometimes underlining the evolution of lineage-specific mechanisms.

The green alga *Chlamydomonas reinhardtii* is a classical reference organism for studying photosynthesis, chloroplast biology, cell cycle control, and cilia structure and function (Salomé and Merchant, 2019). This alga was the first unicellular organism in which miRNAs were described and deep sequencing of short RNAs revealed that it contains a wide variety of endogenous sRNAs (Molnár et al., 2007, Zhao et al., 2007; Voshall et al., 2015; Müller et al., 2020). *Chlamydomonas* also encodes a complex RNA interference (RNAi) machinery consisting of three DCL proteins and three Argonautes but, like most metazoans, lacks a canonical RNA dependent RNA polymerase (Casas-Mollano et al., 2008; Voshall et al., 2015; Valli et al., 2016; Chung et al., 2019). In addition, distinct from land plants, *Chlamydomonas* miRNAs appear to use a metazoan-like seed matching rule to identify their target transcripts but, distinct from animals, the binding sites are predominantly located within the mRNA coding sequences rather than in the 3’ untranslated regions (UTRs) (Yamasaki et al., 2013; Chung et al., 2017; Iwakawa and Tomari, 2017). Nonetheless, the role(s) of most endogenous sRNAs and core RNAi components, particularly with regard to pathway specialization in *Chlamydomonas*, remains to be explored. The highly similar AGO2 and AGO3 proteins (∼90% amino acid identity) bind preferentially to 20-22 nt sRNAs (Voshall et al., 2015; Chung et al., 2019) and the cytoplasmically located AGO3 has been shown to associate with most *bona fide* miRNAs and mediate target transcript cleavage and/or translational repression (Voshall et al., 2015; Yamasaki et al., 2016; Chung et al., 2019). On the other hand, the biological function(s) of AGO1 remains largely uncharacterized. To gain further insights into the evolution of sRNAs and RNAi-related pathways in eukaryotes, particularly in unicellular organisms, we have characterized a novel class of >26 nt small RNAs that associate preferentially with AGO1 in *Chlamydomonas*.

## Results

### AGO1-associated small RNAs

Deep sequencing of total sRNA libraries from vegetative *C. reinhardtii* cells revealed that this unicellular organism contains a complex array of small RNAs, including miRNAs and a variety of endogenous sRNAs (Molnár et al., 2007, Zhao et al., 2007; Voshall et al., 2015; Müller et al., 2020). Intriguingly, more than 36% of the sequenced sRNAs are longer than 26 nts (Figure 1). In contrast, the abundance of >26 nt sRNAs is fairly low in (non-reproductive) whole organism libraries from land plants (Figure 1) (Axtell, 2013; Chávez Montes et al., 2014; Hardcastle et al., 2018; Feng et al., 2020; Lunardon et al., 2020; Chen et al., 2021), including mosses (e.g., *Physcomitrium patens*), liverworts (e.g., *Marchantia polymorpha*), gymnosperms (e.g., *Picea abies* and *Ginkgo biloba*) and angiosperms (e.g., *Arabidopsis thaliana* and *Zea mays*). sRNAs longer than 26 nts are also rare in *Volvox carteri* (Figure 1) (Li et al., 2014; Dueck et al., 2016), a green alga belonging to the same family, Volvocaceae, as *Chlamydomonas* (Craig et al., 2021). We acknowledge that sRNA libraries in the public domain have been generated and sequenced with different techniques and may not be directly comparable but, despite the wealth of published information on sRNAs in land plants (Axtell, 2013; Chávez Montes et al., 2014; Hardcastle et al., 2018; Feng et al., 2020; Lunardon et al., 2020; Chen et al., 2021), we are not aware of any report on the existence of long sRNAs similar to those in *C. reinhardtii*. At present, little is known about the biogenesis and function of these long sRNAs.

**Figure 1.**
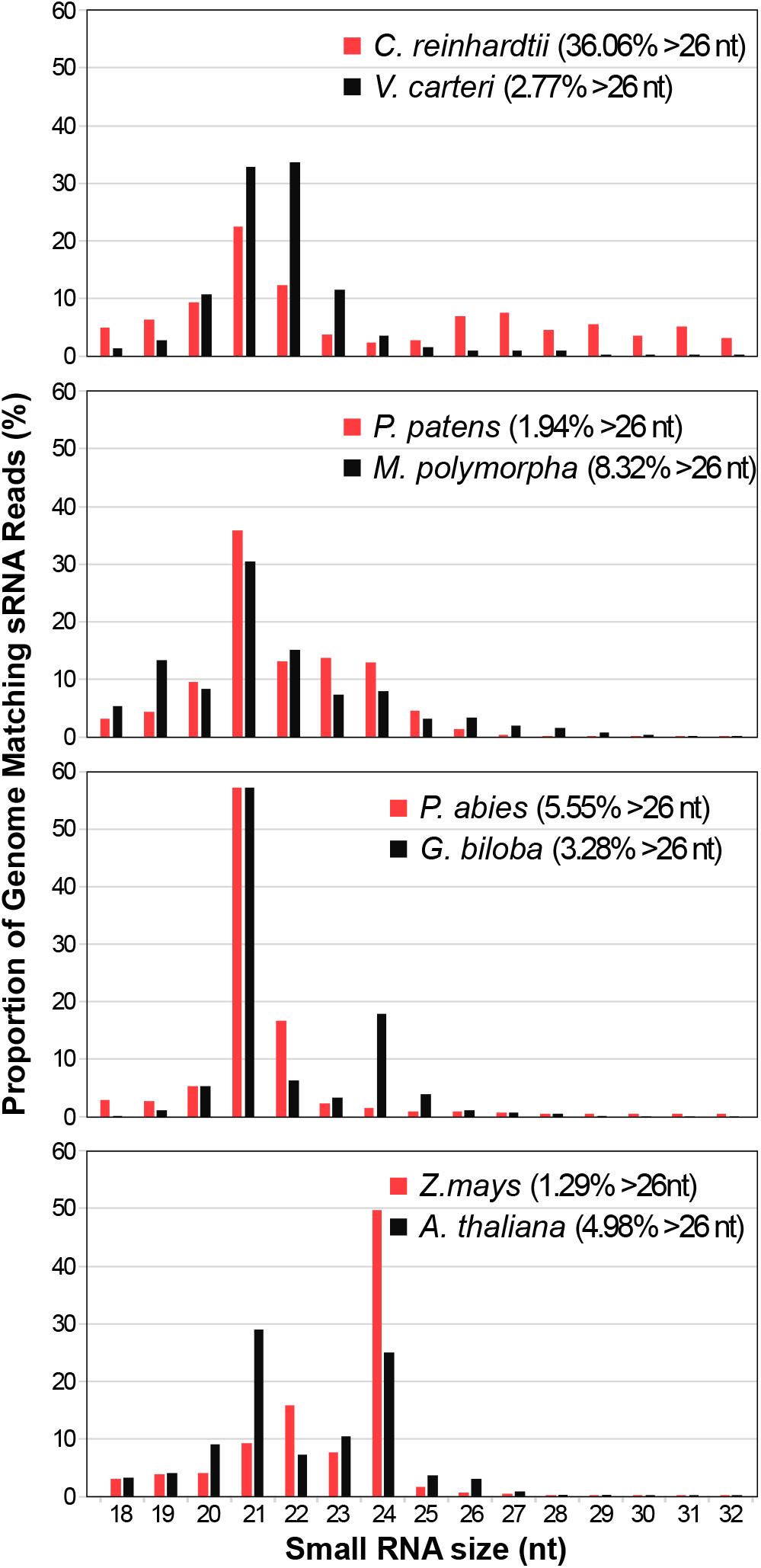
Size distribution of genome mapped total small RNAs in green algae and land plants. nt, nucleotide. Green algal species: *Chlamydomonas reinhardtii* and *Volvox carteri*. Land plant species: *Physcomitrium patens*, *Marchantia polymorpha*, *Picea abies*, *Ginkgo biloba*, *Zea mays* and *Arabidopsis thaliana*.

Similarly, the role(s) of AGO1, the most abundant Argonaute in *Chlamydomonas* (Chung et al., 2019), is not known. Phylogenetic analyses indicate that *C. reinhardtii* AGO1 and AGO2/AGO3 belong to different clades, possibly having diverged in a common ancestor of the Chlorophyceae algal class (Supplemental Figure S1). However, putative orthologs of AGO1 appear to have been lost in some lineages of the Chlorophyceae (including *V. carteri*) (Supplemental Figure S1). From a functional perspective, *Chlamydomonas* AGO1 contains the four conserved domains typical of eukaryotic AGOs, namely N-terminal, PAZ, MID and PIWI domains (Casas-Mollano et al., 2008; Chung et al., 2019). Moreover, the PIWI domain includes the RNase H-like active site (Iwakawa and Tomari, 2022), with putative functional residues (i.e., DDDD) in the catalytic tetrad, suggesting that AGO1 has the capability to cleave target transcripts (Chung et al., 2019).

As previously described for AGO3 (Voshall et al., 2015), we introduced a transgene expressing FLAG-tagged AGO1 into the *Chlamydomonas* Maa7-IR44 strain (Ma et al., 2013). Small RNAs associated with AGO1 were then isolated by co-immunoprecipitation with the FLAG-tagged protein. Deep sequencing of the corresponding sRNA libraries generated 18,791,879 total reads longer than 18 nts, of which 9,195,517 and 1,589,069 mapped perfectly to the nuclear and the chloroplast genomes, respectively. Examination of the genome-matching reads revealed that >75% of the AGO1-associated sRNAs were longer than 26 nts (Figure 2A). By contrast, as already reported (Voshall et al., 2015; Chung et al., 2019), AGO3 associates preferentially with sRNAs 20-22 nt in length and only 0.1% were longer than 26 nts (Figure 2A). Redundant sRNAs associated with AGO1 matched predominantly to introns (∼21.4%) and 3’ UTRs (∼25.7%) of predicted protein coding genes as well as to intergenic regions (∼22.5%) in the nuclear genome (Figure 2B). AGO1 also bound to abundant tRNA fragments derived from genes encoded in the nuclear (∼5.1%) or chloroplast (∼14.9%) genomes (Figure 2B). Less than 0.2% of the AGO1 sRNAs matched to known transposable elements (Figure 2B). The expression of a subset of AGO1-associated long sRNAs was verified by northern blot analyses (Figure 2C), which also demonstrated the existence of ladders of size variants ranging from 26 to almost 42 nts in length. tRNA-derived fragments were instead more precisely defined in size (Figure 2C).

**Figure 2.**
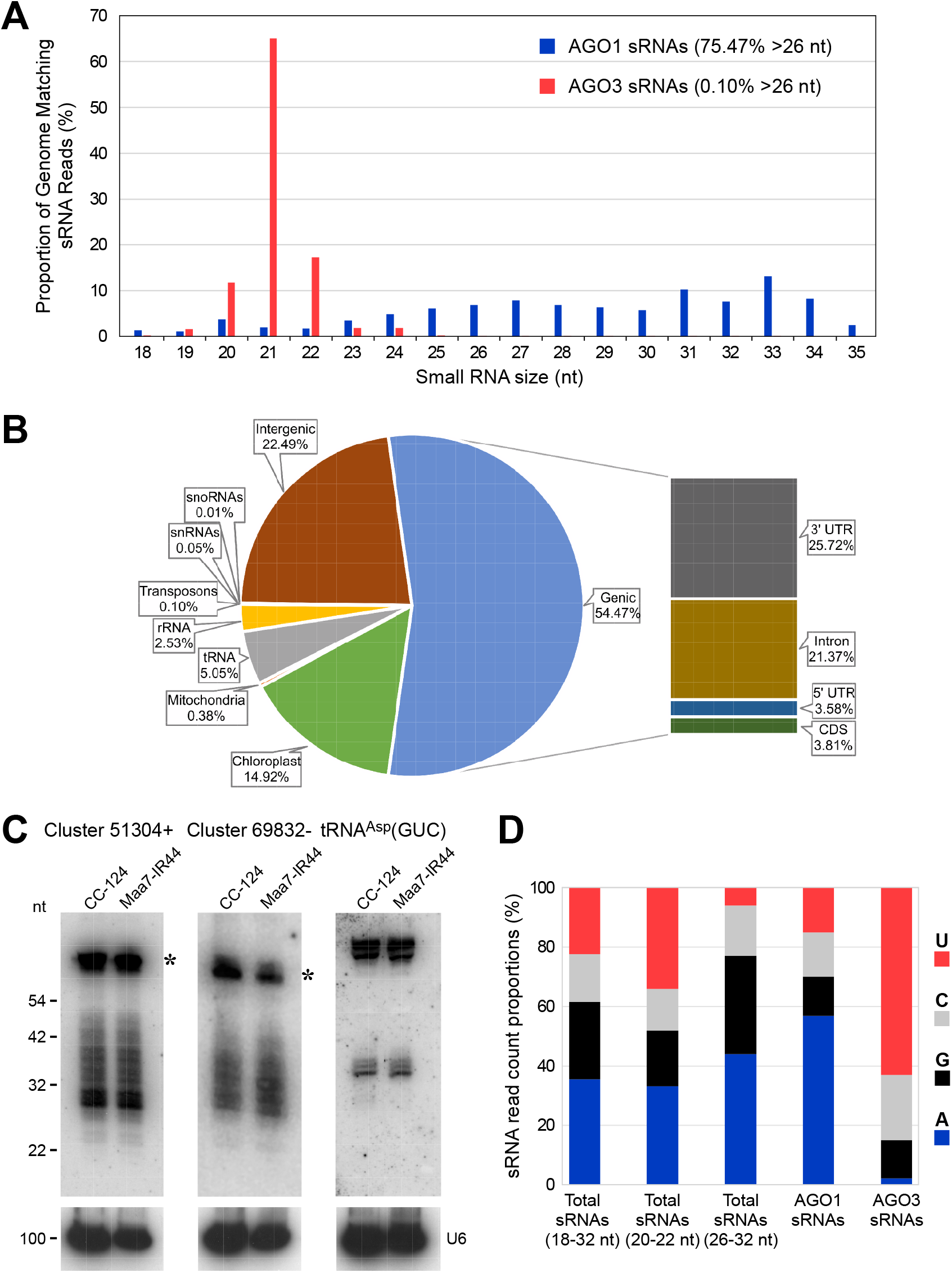
AGO1-associated small RNAs in *Chlamydomonas reinhardtii*. A, Size distribution of genome mapped AGO1-associated sRNAs (blue) and AGO3-associated sRNAs (red). B, Abundance (as % of all genome mapped reads) of AGO1-associated redundant sRNAs matching sequences of distinct functional categories. The chloroplast fraction corresponds to tRNAs encoded in the chloroplast genome. C, Northern blot analyses of small RNAs isolated from the indicated strains and detected with probes specific for the most abundant long sRNAs of Clusters 51304+ or 69832-(Supplemental Table S5) or tRNA fragments derived from the nucleus encoded tRNA^Asp^(GUC). The same filters were reprobed with the U6 small nuclear RNA sequence (U6) as a control for equivalent loading of the lanes. The asterisks indicate putative RNA precursors of the long sRNAs. CC-124, wild type strain; Maa7-IR44, CC-124 transformed with an inverted repeat (IR) transgene designed to induce RNAi of *MAA7* (encoding tryptophan synthase β subunit). D, 5’ terminal nucleotide preference of genome mapped total sRNAs (of the indicated size fractions), AGO1-associated sRNAs and AGO3-associated sRNAs.

We also examined the 5’ terminal nucleotide bias in total sRNA libraries from wild type *C. reinhardtii* strains, which presumably include sRNAs associated with AGO1, AGO2 and AGO3 as well as some unbound sRNAs. Relative to the total sRNA population (i.e., sRNAs of 18-32 nt in length), the fraction 26-32 nt in length was enriched in sRNAs starting with A, whereas the fraction 20-22 nt in length was enriched in sRNAs starting with U (Figure 2D). Consistent with these findings and as an indication of binding specificity, 57% of AGO1-associated sRNAs (which are mainly longer than 26 nts) had an A at the 5’-most position (Figure 2D), whereas the majority of the AGO3 bound sRNAs had a U as the 5’ nucleotide (Voshall et al., 2015).

To corroborate that the *C. reinhardtii* long sRNAs associate with native AGO1, we used CRISPR/Cas9 genome editing (Akella et al., 2021) to generate knockout mutants of the *AGO1* gene. Cells of the wild type CC-124 strain were electroporated with both a CRISPR/Cas9 ribonucleoprotein (RNP) targeting the *AGO1* exon1 and a single-stranded oligodeoxynucleotide (ssODN), with a modified DNA sequence, overlapping the Cas9 cleavage site. Precise repair of the DNA double-strand break caused by the CRISPR/Cas9 RNP, using the ssODN as a homologous template, would substitute 8 base pairs (bp) within the coding sequence of the *AGO1* exon 1 (Figure 3A). This introduces an in-frame stop codon and destroys the Cas9 protospacer adjacent motif. We identified two (independently generated) precisely edited mutants by PCR analyses of selected *Chlamydomonas* colonies, using a primer that anneals exclusively to the altered sequence (Figure 3, A and B). Sequencing of a 235 bp PCR product (Figure 3A, primers F1/R2), overlapping the mutated site, revealed no additional, unintended changes to the *AGO1* exon 1 DNA sequence. In the edited mutants, impaired formation of AGO1 effector complexes is expected to result in destabilization of the associated sRNAs, although some sRNAs may persist as unbound or by alternative association with the intact AGO2/AGO3 proteins. Indeed, northern blot analyses showed that the abundance of all tested long sRNAs was reduced in the *ago1* knockout mutants, in comparison to the wild type strain (Figure 3C and Supplemental Figure S2). The results, taken together, demonstrate that *C. reinhardtii* contains a unique class of long sRNAs, with a bias for adenine as the starting nucleotide, which binds preferentially to AGO1.

**Figure 3.**
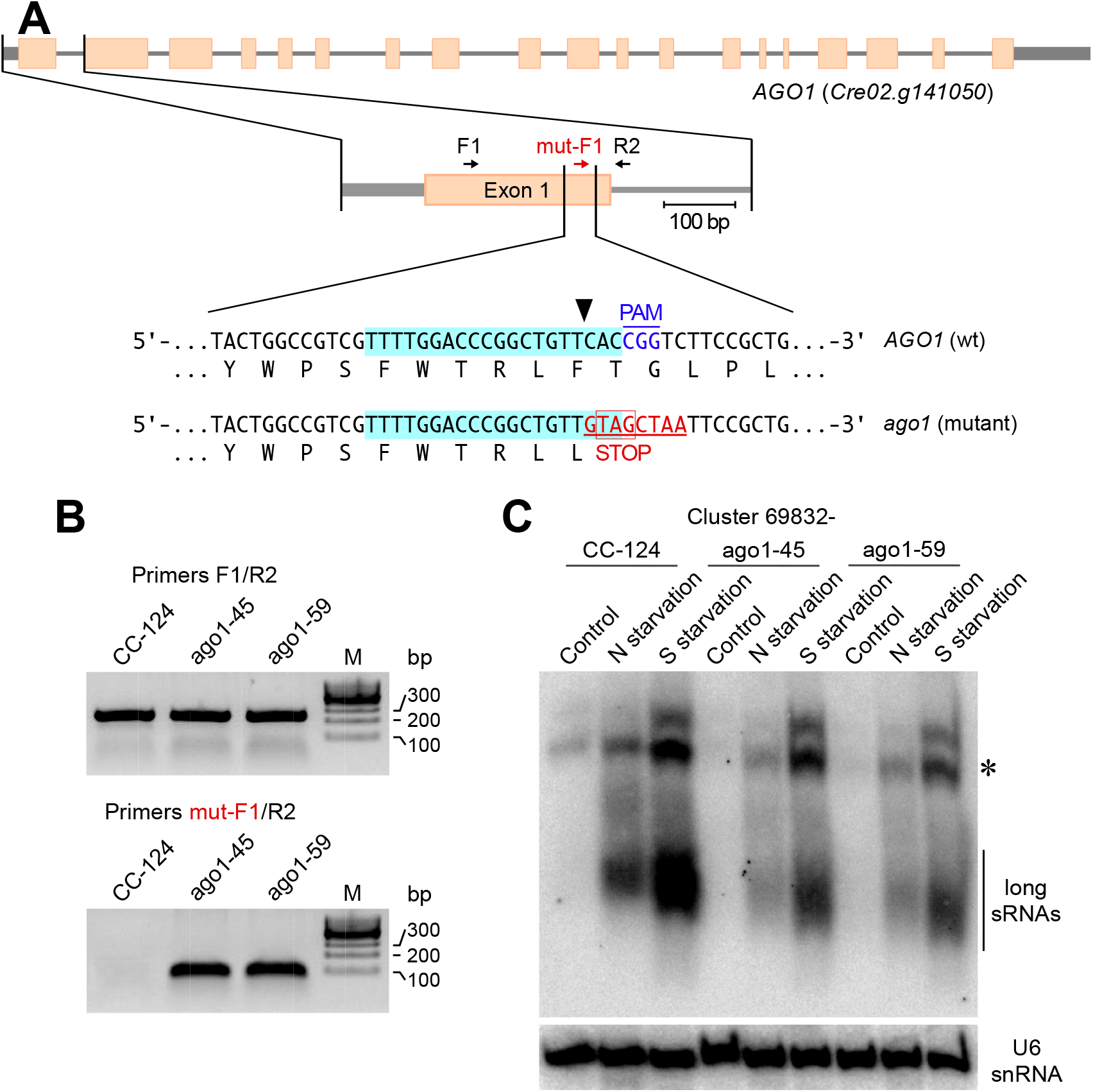
Long sRNAs in *AGO1* edited mutants. A, Schematic of the *AGO1* gene showing the CRISPR/Cas9 target region in exon 1. Short arrows indicate primers used for PCR analyses. In the wild type sequence (top sequence), the target region is highlighted in light blue. The Cas9 cleavage site is indicated by a black arrowhead and the protospacer adjacent motif (PAM) is shown in blue font. Homology directed repair of the DNA double-strand break, using as template the electroporated ssODN, introduces 8 base pair changes (underlined red font) into the genome, including an in-frame stop codon (bottom sequence). B, The *AGO1* target region was amplified by PCR with primers F1/R2 (annealing outside the edited sequence) or mut-F1/R2 (with one primer annealing exclusively to the edited sequence). The panels show representative reverse images of PCR products resolved by agarose gel electrophoresis. CC-124, wild type strain; ago1-45 and ago1-59, edited mutants with a disrupted *AGO1* gene in the CC-124 background. C, Northern blot analysis of long sRNAs from cluster 69832-in the indicated strains cultured in nutrient replete medium (Control) or under nitrogen (N starvation) or sulfur (S starvation) deprivation for 24 h. The same membrane was reprobed with the U6 small nuclear RNA sequence as a control for equivalent loading of the lanes. The asterisk indicates putative RNA precursors of the long sRNAs.

### Genomic loci encoding AGO1-associated long sRNAs

Abundant long sRNAs (excluding tRNA fragments) mapped to a limited number of genomic loci in *C. reinhardtii* chromosomes 1, 3, 6, 7, 8, and 9 (Figure 4A). However, the sequences of most long sRNAs were partly homologous to each other (Table 1 and Supplemental Figure S3), corresponding to moderately repetitive elements. Adjacent reads mapping to the nuclear genome no more than 100 nt apart, regardless of strand, were annotated as belonging to the same long sRNA cluster (Supplemental Figure S3).

**Figure 4.**
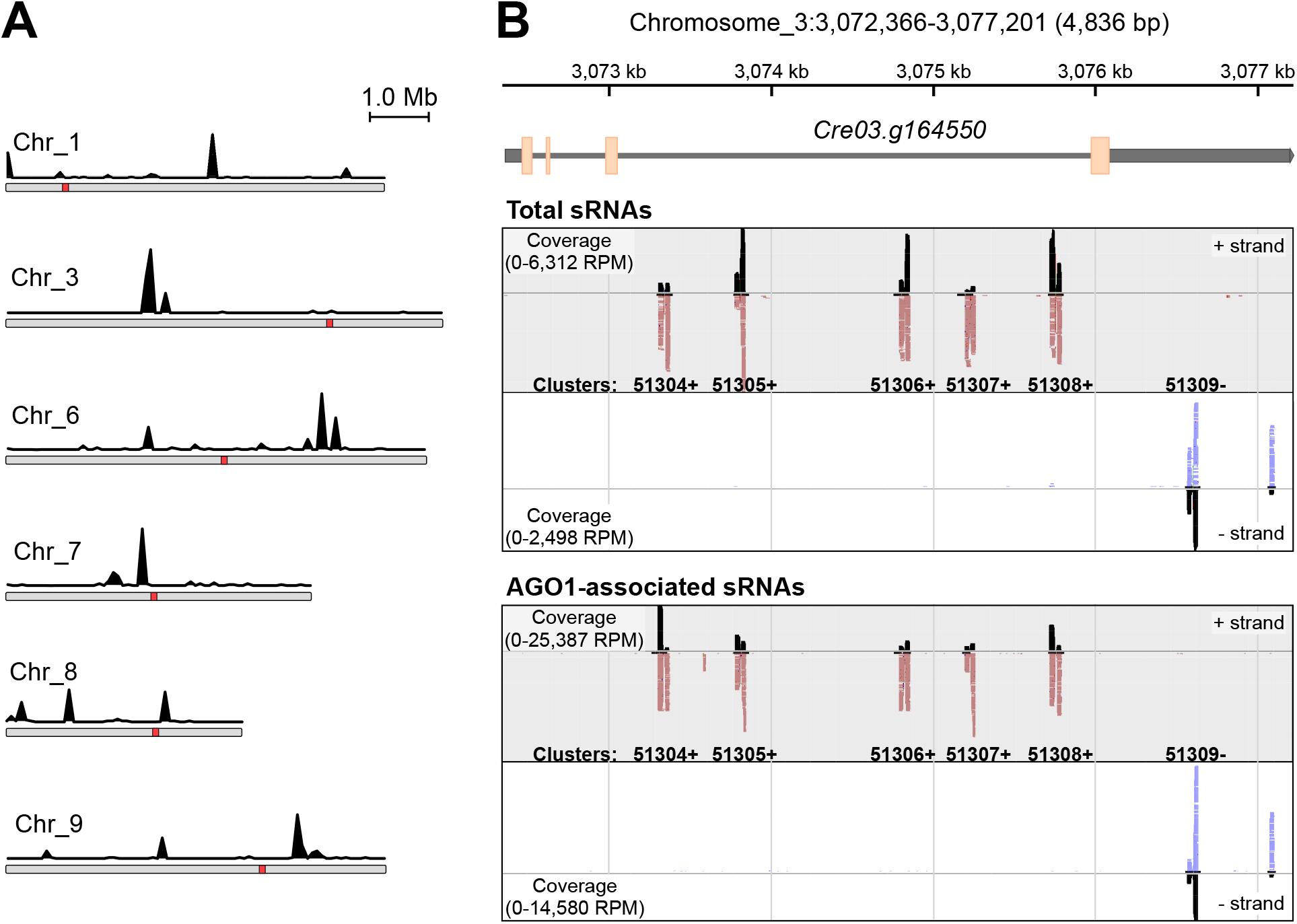
AGO1-associated long sRNAs are encoded in clustered genomic loci. A, Location of genomic loci encoding AGO1-associated long sRNAs on *Chlamydomonas* chromosomes. The black traces indicate the number of AGO1-associated long sRNAs (from normalized libraries) mapped in a sliding 10-kb window across the *C. reinhardtii* chromosomes. Red squares represent putative centromeres (Craig et al., 2021). B, Genome browser view of the clusters matching to the *Cre03.g164550* gene, encoding a homolog of the conserved eukaryotic FRA2/BolA-like protein 2. The upper panel shows the chromosome 3 coordinates and a diagram of the *Cre03.g164550* gene with exons (orange boxes), introns (thin gray lines) and untranslated regions (thick gray lines). The middle panel shows sRNAs from total sRNA libraries mapping to the sense strand (upper) or the antisense strand (lower). Individual mapped sRNAs are shown as red (mapped on + strand) or blue (mapped on – strand) along with coverage in black [indicated as reads per million (RPM)]. The bottom panel shows AGO1-associated long sRNAs mapping to the sense strand (upper) or the antisense strand (lower).

**Table 1.**
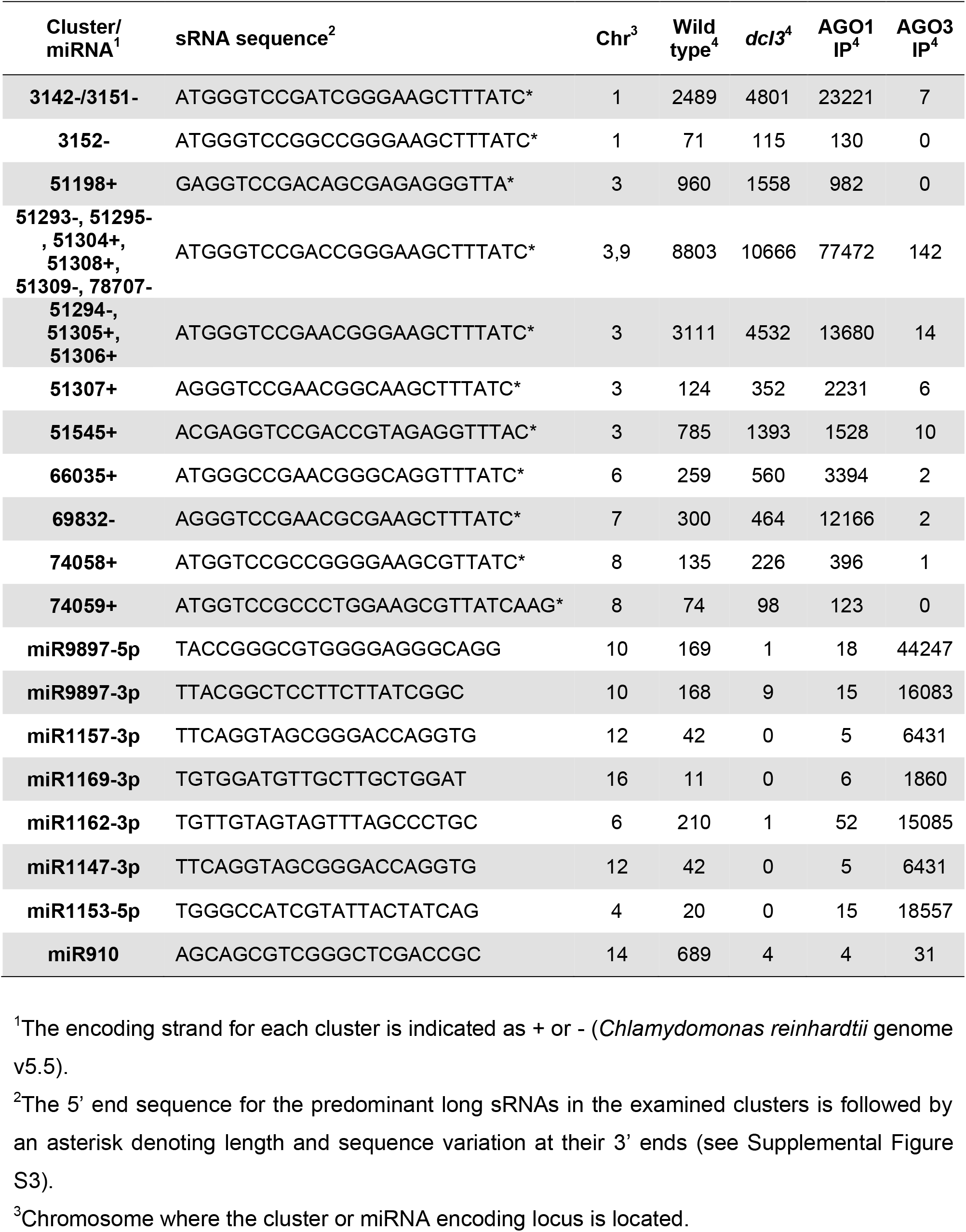
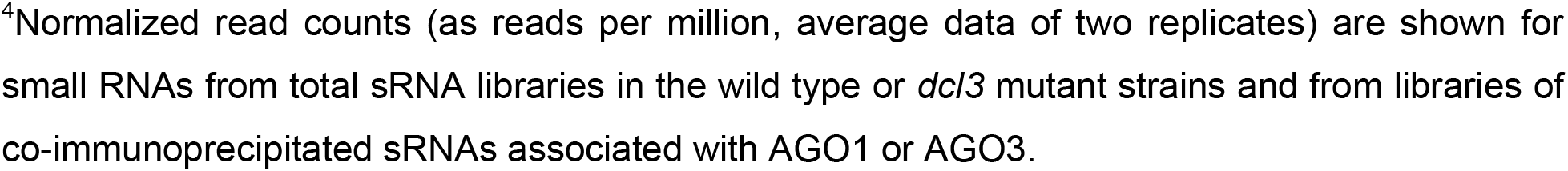
AGO1-associated long sRNAs and miRNAs in *Chlamydomonas reinhardtii*

Additionally, these clusters of long sRNAs encoding sequences tended to form superclusters at certain genomic locations. As an example, the *C. reinhardtii Cre03.g164550* gene encodes a homolog of the conserved eukaryotic FRA2/BolA-like protein 2, which in both fungi and mammals plays an essential role in trafficking [2Fe-2S] clusters to certain enzymes and transcription factors that control iron metabolism (Rey et al., 2019; Talib and Outten, 2021). The third intron of this gene is ∼3 kb in length and contains 5 clusters encoding AGO1-associated long sRNAs in sense orientation (Figure 4B, clusters 51304+, 51305+, 51306+, 51307+ and 51308+). An additional cluster is located in the 3’ UTR of the same gene in antisense orientation (Figure 4B, cluster 51309-). The structure of the *FRA2/BolA-like protein 2* gene, with 4 exons and 3 introns, is conserved in algae of the Trebouxiophyceae (e.g., *Coccomyxa subellipsoidea*) and Chlorophyceae (e.g., *C. reinhardtii*) classes, which diverged 750-850 MYA (Leliaert et al., 2012). However, in most algal species the third intron is relatively short including, for instance, *Tetradesmus obliquus*, *V. carteri* and *Edaphochlamys debaryana* within the Chlorophyceae (Supplemental Figure S4). In contrast, this third intron has experienced substantial size expansion in *C. reinhardtii* and its close relatives, *Chlamydomonas incerta* and *Chlamydomonas schloesseri* (Supplemental Figure S4), when compared with the more distant relative *E. debaryana* [for details on the phylogenetic relationships see Craig et al. (2021)]. Interestingly, the three *Chlamydomonas* species contain, in the third intron and 3’ UTR of their *FRA2/BolA-like protein 2* genes, conserved sequences homologous to the AGO1-associated long sRNAs (Figure 4B and Supplemental Figures S5 and S6); and these sequences have been amplified so that the size of the third intron has increased in parallel with the number of long sRNA encoding clusters (1 in *C. shloesseri*, 3 in *C. incerta*, and 5 in *C. reinhardtii* at a threshold of 70% nucleotide identity).

We also examined expression, by northern blot analyses, of sequences homologous to clusters 51304+, 51545+ and 69832-in *E. debaryana*, *C. incerta*, and *C. reinhardtii* (Supplemental Figure S7, A, C, and D). Using probes specific for the most abundant reads of each of those clusters, long sRNAs were detected in *C. reinhardtii* and its close relative *C. incerta*. Moreover, changes in steady state levels of the related long sRNAs, in response to several nutrient deprivation conditions (see below), were conserved in these two species (Supplemental Figure S7, A, C, and D). Reprobing the same membrane with an oligonucleotide hybridizing to a *C. reinhardtii* miRNA, miR912 (21 nt in length), substantiated that the detected sRNAs were indeed larger in size than canonical miRNAs and also showed that miR912 is not conserved in *C. incerta* (Supplemental Figure S7B). The *E. debaryana* genome does not seem to encode loci with homology to the examined *C. reinhardtii* long sRNAs and we did not observe obvious hybridization to RNA isolated from this species (Supplemental Figure S7). Thus, although the origin of the repetitive genomic loci encoding long sRNAs remains unknown, at least some of these sequences have been conserved and amplified within core *Chlamydomonas* species. Moreover, changes in expression under various nutritional conditions also appear to have been conserved.

### Biogenesis of AGO1-associated long sRNAs

To assess whether the long sRNAs show any evidence of specific processing, we aligned the sRNAs within each cluster to the corresponding genomic sequences and evaluated the precision with which their 5’ and 3’ ends had been generated as well as the possible folding of precursor RNAs into secondary structures. In most cases, a putative precursor with fold-back hairpin secondary structure, ∼60-80 nt in length, could be predicted by using RNAfold from the Vienna RNA package (Lorenz et al., 2011) (Figure 5, cluster 51304+, and Supplemental Figure S3). Moreover, northern blot analyses suggested the existence of easily detectable RNA precursors of the predicted length (Figures 2C and 3C, asterisks). Intriguingly, at least some of these putative RNA precursors also appear to be destabilized in the absence of AGO1 (Figure 3C and Supplemental Figure S2A).

**Figure 5.**
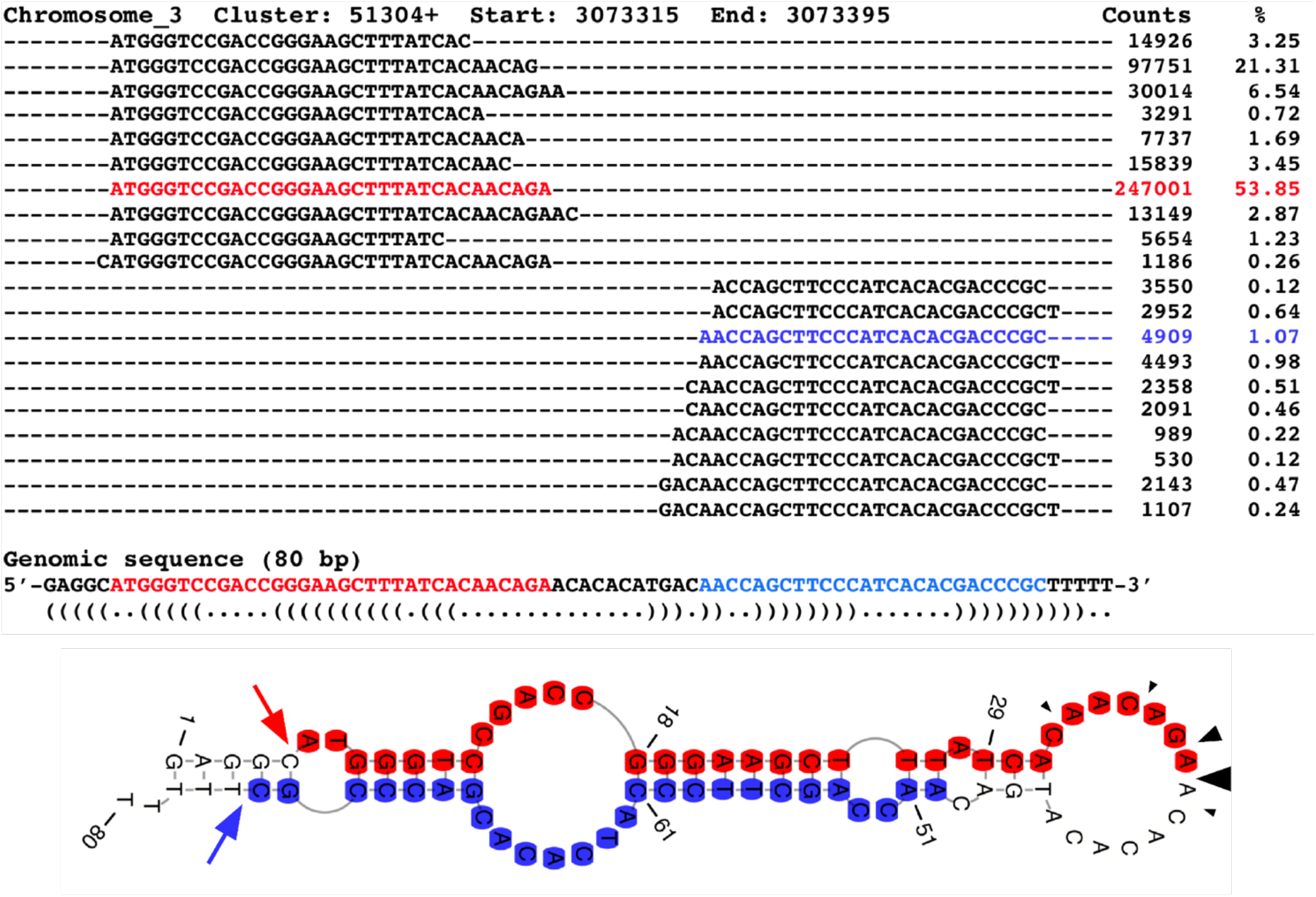
Predicted precursor RNA structure for cluster 51304+. Sequences of AGO1-associated long sRNAs were aligned to the corresponding genomic sequence and their frequency in the libraries is shown as raw counts. The secondary structure of the encoding genomic sequence was predicted with RNAfold (Lorenz et al., 2011) and depicted at the bottom. The most abundant long sRNAs encoded in the 5’ arm or the 3’ arm of the predicted hairpin are indicated in red and blue, respectively. Fairly precise putative cleavage sites on the left side of the depicted hairpin are indicated by red and blue arrows. More imprecise putative cleavage sites in the terminal loop of the depicted hairpin are indicated by black arrowheads.

The 5’ end of the long sRNAs encoded in the 5’ arm of the predicted hairpin and the 3’ end of the long sRNAs encoded in the 3’ arm of the hairpin appeared to be much more precisely processed (Figure 5 and Supplemental Figure S3). This corresponds to both strand ends on the left side of the secondary structure diagrams (Figure 5 and Supplemental Figure S3). However, in very few instances the sequences seemed to form a stable dsRNA stem at the putative cleavage sites (Supplemental Figure S3), making it unlikely that they were processed by RNase III enzymes such as the DCL proteins (Vermeulen et al., 2005; Macrae et al., 2006; Song and Rossi, 2017). Additionally, the processing of the right ends of the hairpin secondary structures (i.e., 3’ end of long sRNAs encoded in the 5’ arm of the hairpin and 5’ end of long sRNAs encoded in the 3’ arm of the hairpin) appeared to be much more imprecise and the putative cleavage site(s) often corresponded to predicted ssRNA regions. Also, we cannot rule out that some of the apparent specificity in processing 5’ ends of long sRNAs may reflect the predominant stabilization of sequences starting with adenine because of the AGO1 binding preference.

Although the secondary structure of predicted long sRNA precursors suggested that they were not processed by DCL proteins, we tested more directly whether the biogenesis of long sRNAs was dependent on these RNase III endonucleases. As already mentioned, *C. reinhardtii* encodes three DCL proteins, of which DCL2 and DCL3 are fairly similar to each other (Casas-Mollano et al., 2008; Valli et al., 2016). For our studies, we focused on the more divergent DCL1 and DCL3, which are expressed at comparable levels and much more abundantly than DCL2 based on transcript steady-state amounts (Casas-Mollano et al., 2008; Zones et al., 2015). Examination of small RNA libraries from a *C. reinhardtii dcl3* null mutant (Valli et al., 2016) revealed that lack of DCL3 did not affect production of the long sRNAs (>26 nts in length) preferentially associated with AGO1 (Figure 6A and Table 1). However, as already described (Valli et al., 2016), the accumulation of 20-22 nt sRNA (including most miRNAs) is substantially reduced in this mutant (Figure 6A and Table 1).

**Figure 6.**
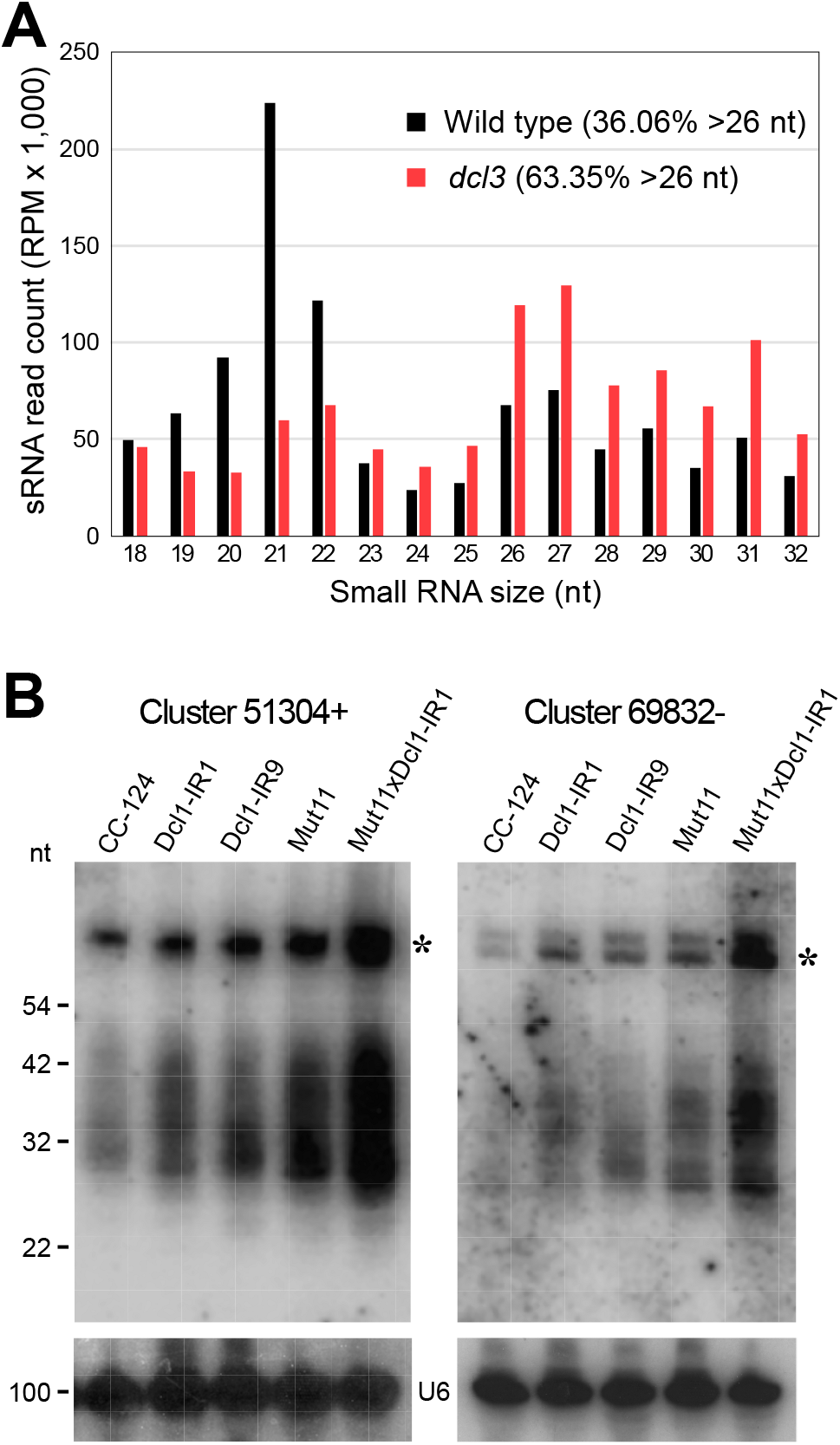
The biogenesis of AGO1-associated long sRNAs is independent of DCL1 and DCL3. A, Size distribution of genome mapped total sRNAs from a wild type strain (black) and a *dcl3* null mutant (red). RPM, reads per million. B, Northern blot analyses of small RNAs isolated from the indicated strains and detected with probes specific for the most abundant long sRNAs of Clusters 51304+ or 69832-(Supplemental Table S5). The same filters were reprobed with the U6 small nuclear RNA sequence (U6) as a control for equivalent loading of the lanes. The asterisks indicate putative RNA precursors of the long sRNAs. CC-124, wild type strain; Dcl1-IR1 or Dcl1-IR9, CC-124 transformed with an IR transgene designed to induce RNAi of *DCL1* (Casas-Mollano et al., 2008); Mut11, mutant defective in a core subunit of H3K4 methyltransferase complexes required for chromatin-mediated silencing (van Dijk et al., 2005); Mut11xDcl1-IR1, Mut11 transformed with an IR transgene designed to induce RNAi of *DCL1* (Casas-Mollano et al., 2008).

We have previously generated *C. reinhardtii* strains where DCL1 expression was suppressed by RNAi in a wild type background (CC-124) and in a mutant (Mut11) defective in chromatin-mediated transcriptional silencing (Casas-Mollano et al., 2008). These strains were used to demonstrate that DCL1 plays a role in transposon repression, particularly when chromatin-mediated silencing was compromised (Casas-Mollano et al., 2008). Northern blot analyses of the same strains indicated that production of the tested AGO1-associated long sRNAs is not affected by DCL1 suppression (Figure 6B). On the contrary, the abundance of long sRNAs and of their putative precursors increases in the DCL1 RNAi lines, and even more so in the strain that is also deficient in chromatin-mediated transcriptional silencing (Figure 6B). Thus, our results strongly suggest that the biogenesis of long sRNAs is independent of DCL1 and DCL3. Moreover, compensatory functions between DCL1 and DCL3 seem unlikely since DCL1 appears to antagonize the production of long sRNAs and it would not restore any hypothetical, DCL3-dependent, processing of long sRNAs in the *dcl3* mutant.

We also attempted to examine mutants of the third DCL protein encoded in the *C. reinhardtii* genome, namely DCL2. However, two putative mutants, with mapped disruptions in the *DCL2* coding sequence in a genome wide insertional mutagenesis library (LMJ.RY0402.066980 and LMJ.RY0402.093816; Li et al., 2019), proved to be incorrect upon detailed genotyping by PCR. Nonetheless, DCL2 is expressed at low levels (Casas-Mollano et al., 2008; Zones et al., 2015), with a protein abundance ∼28-fold lower than that of DCL3 (Supplemental Figure S8A). RT-PCR analyses revealed that its transcript is present at similar levels in the wild type, *dcl3*, and *DCL1* RNAi strains (Supplemental Figure S8B), implying no obvious compensatory up-regulation of *DCL2* expression in mutants or epi-mutants defective in the other DCL proteins. Moreover, DCL2 does not compensate for the lack of DCL3, despite their sequence similarity, in the processing of miRNAs and other 20-22 nt sRNAs in the *dcl3* mutant (Valli et al., 2016). Under nutrient deprivation conditions, when the abundance of long sRNAs increases prominently (see below), we observed an increase in *AGO1* transcript levels but *DCL2* expression did not change significantly (Supplemental Figure S8C). On the whole, it seems unlikely that any *C. reinhardtii* DCL protein might play a major role in generating the AGO1 associated long sRNAs, especially since the predicted precursor RNAs do not appear to be cleaved at dsRNA regions (Supplemental Figure S3). Our observations suggest that the *Chlamydomonas* long sRNAs have a distinct biogenesis from that of canonical miRNAs and siRNAs although the actual molecular mechanism(s) remains elusive.

### Potential role(s) of AGO1-associated long sRNAs

Patterns of adaptive evolution have been observed in animal RNAi-pathway genes involved in the control of transposable elements and/or viruses, such as those coding for PIWI proteins and a subset of insect AGOs including orthologs of *Drosophila melanogaster* AGO2 (Wynant et al., 2017; Palmer et al., 2018; Ozata et al., 2019). Their accelerated evolution has been hypothesized to reflect an ‘evolutionary arms race’ with the rapidly evolving molecular parasites they are targeting (Wynant et al., 2017; Palmer et al., 2018). Directional selective pressure can be identified by testing for an elevated rate of nonsynonymous nucleotide substitutions (Ka) relative to synonymous substitutions (Ks) within the coding sequence of a protein (Hurst, 2002; Wynant et al., 2017). Using tree-based methods for the calculation of Ka/Ks ratios of *Chlamydomonas* AGO genes, substantially elevated values were observed for the *AGO1* and *AGO2/AGO3* genes in comparison with the housekeeping *ACT1* (encoding actin) gene (Supplemental Figure S9). The Ka/Ks ratios for *Chlamydomonas* AGOs, particularly for AGO1, were comparable to those of the fast-evolving animal RNAi-pathway genes (Wynant et al., 2017), suggesting that AGO1 might likewise play a role in defense responses against transposons and/or viruses.

As described above, AGO1 associates predominantly with uncommonly long sRNAs derived from moderately repetitive sequences and, in principle, it might also have a role in regulating the expression of endogenous genes using the associated long sRNAs as guides. To obtain insight into the possible function(s) of these long sRNAs, we examined the expression, under a variety of stress conditions, of the abundant sequences corresponding to cluster 51304+. Interestingly, in northern blot analyses of *C. reinhardtii*, the levels of putative precursor and mature long sRNAs from cluster 51304+ increased markedly under sulfur or nitrogen deprivation, in comparison to those in cells grown in nutrient replete medium (Figure 7A). A similar response was observed in both *C. reinhardtii* and its relative *C. incerta* for long sRNAs corresponding to clusters 51304+, 51545+ and 69832-(Supplemental Figure S7, A, C, and D). Examination of total sRNA libraries generated from *C. reinhardtii* cells cultured under nutrient replete or nitrogen deprived conditions also revealed increased abundance of a subset of the AGO1-associated long sRNAs when nitrogen was lacking in the medium (Supplemental Table S1, clusters 51198+, 51293-, 51295-, 51304+, 51308+, 51309+, 78707-, and 66035+).

**Figure 7.**
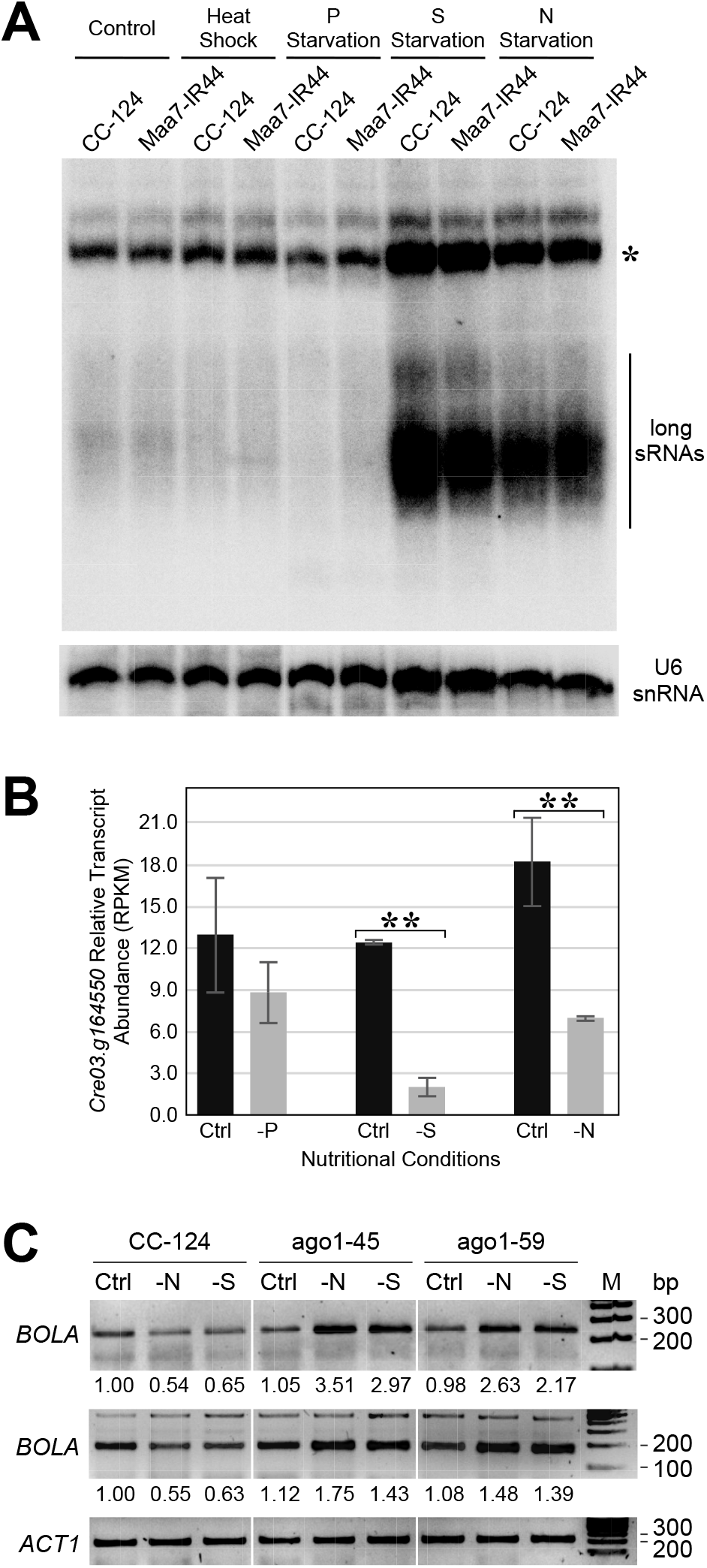
The abundance of AGO1-associated long sRNAs increases under certain nutritional deprivation conditions. A, Northern blot analysis of long sRNAs from cluster 51304+ in *Chlamydomonas* cells cultured under the denoted conditions. Control (Ctrl), nutrient-replete standard laboratory conditions; Heat shock, 42°C for 20 min; P starvation, phosphorus deprivation for 24 h; S starvation, sulfur deprivation for 24 h; N starvation, nitrogen deprivation for 24 h. sRNAs were detected with a probe specific for the most abundant long sRNAs of cluster 51304+ (Supplemental Table S5). The same filter was reprobed with the U6 small nuclear RNA sequence as a control for equivalent loading of the lanes. The asterisk indicates putative RNA precursors of the long sRNAs. CC-124, wild type strain; Maa7-IR44, CC-124 transformed with an IR transgene designed to induce RNAi of *MAA7* (encoding tryptophan synthase β subunit). B, Relative mRNA levels of the predicted long sRNA target gene *Cre03.g164550*, encoding a homolog of the conserved eukaryotic FRA2/BolA-like protein 2, in wild type *C. reinhardtii* grown under the indicated nutritional conditions. RPKM, reads per kilobase per million determined from RNA-seq data. Double asterisks indicate significant differences between the indicated samples p<5×10^-5^. C, *BOLA-like* (*Cre03.g164550*) transcript abundance examined by RT-PCR in cells grown under the indicated conditions. Amplification of the *ACT1* (encoding actin) transcript was used for normalization purposes. The panels show representative reverse images of agarose resolved RT-PCR products stained with ethidium bromide. PCR was carried out with primers BOLA-like-F2/BOLA-like-R (top panel) or BOLA-like-F/BOLA-like-R (middle panel) (Supplemental Table S5). The numbers below the panels indicate relative *BOLA-like* (*Cre03.g164550*) mRNA abundance normalized to *ACT1* transcript abundance. ago1-45 and ago1-59, edited mutants with a disrupted *AGO1* gene in the CC-124 background.

In human AGO2, a conserved helix 7 and residue I365 have been reported to shape the miRNA seed region for rapid target RNA recognition (Klum et al., 2018; Gebert and MacRae, 2019). These features (particularly the equivalent isoleucine and nearby residues) are conserved in *Chlamydomonas* AGO1 (Chung et al., 2019), consistent with a similar guide targeting mode as canonical Argonaute proteins. Thus, we used the psRNATarget web server (Dai et al., 2018), with near default parameters (i.e., following miRNA-like pairing rules), to identify target mRNAs hybridizing to the first 21 nucleotides of the abundant long sRNAs derived from the 5’ arm of cluster 51304+ (Figure 5). We note that this sequence is conserved in several additional long sRNA clusters encoded by the *C. reinhardtii* genome (Table 1 and Supplemental Figure S3, clusters 51293-, 51295-, 51304+, 51308+, 51309+, and 78707-) as well as in genomic loci of the closely related species *C. incerta* and *C. schloesseri*. Thirty-three putative targets were predicted using this approach (Supplemental Table S2), including *Cre03.g164550*, which encodes the 51304+ cluster within its third intron, but also contains complementary sequences (in antisense orientation) within its 3’ UTR (Supplemental Figure S10). The predicted targets appear to participate in a wide variety of cellular functions rather than being confined to specific cellular or metabolic pathways (Supplemental Table S2).

We then examined publicly available RNA-seq data collected from wild type *C. reinhardtii* grown under nutrient replete or under phosphorus-, sulfur- or nitrogen-deprivation conditions, to assess the expression level of the predicted targets of the cluster 51304+ long sRNAs (Supplemental Table S3). Of note, *Cre03.g164550* transcript abundance was significantly downregulated under sulfur or nitrogen deprivation (Figure 7B and Supplemental Table S3), coincidental with increased expression of the cluster 51304+ long sRNAs (Figure 7A). Conversely, in the *ago1* edited mutants, which showed under all examined trophic conditions diminished steady-state levels of long sRNAs relative to the wild type strain (Figure 3C and Supplemental Figure S2), the abundance of the *Cre03.g164550* transcript increased under sulfur or nitrogen deprivation (Figure 7C). With the caveat that some target predictions may represent false positives or the predicted binding sites may not be accessible for interaction with a guide sRNA, in the wild type strain 45.5% of the predicted target transcripts were downregulated under sulfur deprivation and 42.4% of the predicted target transcripts were downregulated under nitrogen deprivation (Supplemental Table S3). These observations strongly suggest that AGO1-associated long sRNAs may indeed target specific genes for silencing under certain nutritional conditions.

## Discussion

Noncoding RNAs constitute a sizeable fraction of eukaryotic transcriptomes and some play essential biological roles (Amaral et al., 2008; Palazzo and Koonin, 2020; Tsuzuki et al., 2020). RNA interference, mediated by small noncoding RNAs, is a highly-conserved process influencing gene regulation, genome stability as well as defense responses against genomic parasites (Ghildiyal and Zamore, 2009; Borges and Martienssen, 2015; Wendte and Pikaard, 2017; Yu et al., 2017; Bartel, 2018; Lee and Carroll, 2018; Ozata el al., 2019; Chen and Rechavi, 2021). In multicellular eukaryotes, such as land plants, animals and fungi, extensive duplication and diversification of RNAi machinery components have resulted in complex, partly overlapping pathways for epigenetic regulation (Ghildiyal and Zamore, 2009; Czech and Hannon, 2011; Borges and Martienssen, 2015; Lee and Carroll, 2018; Wang et al., 2021). Duplicated RNAi components appear to allow the evolution of new gene regulatory mechanisms that use distinct sRNAs as sequence-specific determinants, as well as more effective strategies to counteract the action of invading viruses and transposable elements. In contrast, many unicellular eukaryotes seem to have lost entirely the RNAi machinery or have retained only a basic set of RNAi components (Cerutti and Casas-Mollano, 2006; Casas-Mollano et al., 2008). The alga *Chlamydomonas reinhardtii* has long been used as a model system for the characterization of plant processes, such as photosynthesis, and of certain animal structures, such as cilia (Salomé and Merchant, 2019). Intriguingly, *Chlamydomonas* has an RNAi machinery much more complex than might be expected for a unicellular organism, including three AGOs, three DCLs and a diverse assortment of sRNAs (Molnár et al., 2007, Zhao et al., 2007; Casas_Mollano et al., 2008; Voshall et al., 2015; Valli et al., 2016; Chung et al., 2019; Müller et al., 2020). Additionally, *Chlamydomonas* miRNAs do not seem to regulate endogenous mRNA targets through the plant model of high complementarity between miRNA and transcript (Yamasaki et al., 2013; Chung et al., 2017; Iwakawa and Tomari, 2017). Of the core RNAi proteins, *Chlamydomonas* DCL3 and AGO3 have been implicated in the biogenesis and activity of many (all) miRNAs (Voshall et al., 2015; Valli et al., 2016; Yamasaki et al., 2016; Chung et al., 2019). Their closely related paralogs, DCL2 and AGO2, are expressed at low levels (Zones et al., 2015) and AGO2 binds exclusively to ∼21 nt siRNAs of unknown function (Chung et al., 2019). In contrast, the more divergent paralogs DCL1 and AGO1 have not been characterized in detail (Casas-Mollano et al., 2008). Moreover, the biogenesis and function(s) of most endogenous sRNAs in *C. reinhardtii*, including a unique, abundant class of >26 nt sRNAs (Figure 1), remain to be elucidated.

As described here, AGO1 associates predominantly with these long sRNAs, having a bias for adenine as the 5’ terminal nucleotide (Figures 2 and 3). Abundant long sRNAs are generated from a limited number of genomic loci (Figure 4 and Table 1), corresponding to moderately repetitive sequences that match either intergenic regions or introns/3’ UTRs of predicted protein coding genes (Figure 2B). These features are in agreement with several characteristics of the recently annotated locus class 4, distinguished by using data-driven machine learning approaches in a systematic classification of sRNA loci in *C. reinhardtii* (Müller et al., 2020). Moreover, long sRNA encoding sequences, as well as their expression, have been conserved in phylogenetically related *Chlamydomonas* species (Supplemental Figures S4 to S7). The long sRNAs are likely processed from ∼60-80-nt precursor RNAs that can fold into hairpin secondary structures (Figure 5 and Supplemental Figure S3). However, the majority of the putative precursor RNAs do not appear to be cleaved at dsRNA regions (Supplemental Figure S3) and long sRNA biogenesis is independent of DCL1 and DCL3 (Figure 6 and Table 1). Since the putative precursor RNAs seem to be destabilized in the absence of AGO1 (Figure 3C and Supplemental Figure S2A), we speculate that their partial dsRNA nature may facilitate loading into AGO1 effector complexes but elucidating the actual processing mechanism will require additional work.

The *Chlamydomonas* long sRNAs share some features with animal piRNAs (Ghildiyal and Zamore, 2009; Ozata et al., 2019) such as a longer size than canonical miRNAs/siRNAs, being encoded in genomic superclusters and, likely, Dicer-independent biogenesis. However, piRNAs are processed from ssRNAs without obvious secondary structures and they associate with PIWI proteins (Ghildiyal and Zamore, 2009; Ozata et al., 2019), which are absent from plant and green algal species (Cerutti and Casas-Mollano, 2006; Swarts et al., 2014). In land plants, most sRNAs are shorter than ∼26 nts (Axtell, 2013; Chávez Montes et al., 2014; Borges and Martienssen, 2015; Hardcastle et al., 2018; Feng et al., 2020; Lunardon et al., 2020; Chen et al., 2021) and, to our knowledge, sRNAs with similar characteristics to the *Chlamydomonas* long sRNAs have not been described. Thus, the *Chlamydomonas* long sRNAs appear to add to a growing list of unorthodox sRNAs, with unique biogenesis pathways and/or functions (Rzeszutek and Betlej, 2020; Wu et al., 2020; Alves and Nogueira, 2021; Meseguer, 2021).

Based on its fairly high Ka/Ks ratio (Supplemental Figure S9), similar to that of PIWIs and certain animal Argonautes involved in the control of viruses and/or transposons (Wynant et al., 2017; Palmer et al., 2018; Ozata et al., 2019), we hypothesize that *Chlamydomonas* AGO1 may play a role in defense responses against molecular parasites. However, viruses affecting *Chlamydomonas* have not been characterized and sequences matching known transposable elements represent only a minor fraction of the sRNAs associated with AGO1 (Figure 2B). Nonetheless, as previously described (Jeong et al., 2002; Casas-Mollano et al., 2008; Kim et al., 2015), multiple, partly redundant epigenetic mechanisms are involved in preventing transposon mobilization in *Chlamydomonas*. In cells grown under standard laboratory conditions, transposons appear to be primarily repressed by a chromatin-based mechanism(s) and only when this system is defective the involvement of RNAi as a transposon silencing mechanism can be fully appreciated (Jeong et al., 2002; Casas-Mollano et al., 2008; Kim et al., 2015).

AGO1 may also have a role in regulating the expression of certain endogenous genes using the associated long sRNAs as guides. Interestingly, we observed that the abundance of several AGO1-associated long sRNAs is markedly enhanced under certain nutritional stresses, such as nitrogen or sulfur deprivation (Figure 7A, Supplemental Figure S7 and Supplemental Table S1). Under these same conditions, a substantial fraction of 33 putative target transcripts, identified by sequence complementarity to the most abundant reads from cluster 51304+, showed a reduction in mRNA levels (Supplemental Table S3). One of the downregulated transcripts corresponded to the *FRA2/BolA-like protein 2* gene (Figure 7B), which hosts several long sRNA clusters within its third intron but also contains complementary target sequences within its 3’ UTR (Figure 4B and Supplemental Figure S10). In contrast, in the *ago1* edited mutants, the transcript abundance of the *FRA2/BolA-like protein 2* gene increased under sulfur or nitrogen deprivation (Figure 7C). These observations, taken together, are consistent with target transcript decay triggered by an effector complex comprised of AGO1 and its associated long sRNAs.

Curiously, some of the conserved cluster sequences encoding long sRNAs, at least those within the third intron of the *FRA2/BolA-like protein 2* gene, have been amplified in closely related *Chlamydomonas* species (Supplemental Figures S4 to S6) but they do not match known transposable elements. However, both DCL1 and MUT11, which play a role in controlling transposons (Jeong et al., 2002; van Dijk et al., 2005; Casas-Mollano et al., 2008), appear to antagonize the production of long sRNAs and of their putative RNA precursors (Figure 6B). Thus, although the origin of the long sRNA encoding sequences remains elusive, in several respects they appear to have transposon-like behavior. We surmise that the repetitive sequences encoding long sRNAs might have been ancestrally targeted for silencing by AGO1 effector complexes. Yet, in the course of evolution, some of the long sRNAs may have fortuitously acquired endogenous target genes, conferring a selective advantage in gene regulation and leading to fixation of the encoding clusters.

At their core, eukaryotic RNAi systems consist of a short guide RNA, allowing for sequence-specific target recognition, and an effector protein of the Argonaute family, which mediates downstream effects with varying outcomes (Burroughs et al., 2014; Swarts et al., 2014; Dexheimer and Cochella, 2020). The guide sRNAs identified in earlier studies were derived from dsRNA precursors processed by RNAse III type Dicer endonucleases (Ghildiyal and Zamore, 2009; Burroughs et al., 2014; Borges and Martienssen, 2015; Yu et al., 2017; Bartel, 2018; Chen and Rechavi, 2021). However, it is now becoming apparent that guide sRNAs can be generated by a variety of alternative pathways (Ghildiyal and Zamore, 2009; Yang and Lai, 2010; Burroughs et al., 2014; Wendte and Pikaard, 2017; Bartel, 2018; Alves and Nogueira, 2021). The functional role of AGOs also appears to have changed during evolution, from relatively simple host-defense proteins against viruses and transposons to key players in complex multiprotein regulatory pathways in multicellular organisms (Shabalina and Koonin, 2008; Ghildiyal and Zamore, 2009; Burroughs et al., 2014; Swarts et al., 2014; Borges and Martienssen, 2015; Lee and Carroll, 2018; Dexheimer and Cochella, 2020; Chen and Rechavi, 2021). Within this context, the long (>26 nt) sRNAs in the alga *Chlamydomonas reinhardtii* might derive from transposon-like sequences which, during evolution, acquired gene regulatory functions.

## Material and methods

### *C. reinhardtii* strains, mutants, and culture conditions

Strain Maa7-IR44, containing an inverted repeat transgene designed to silence the *MAA7* gene (encoding Tryptophan Synthase β subunit), has been previously described (Ma et al., 2013). Mut11 is an insertional mutant defective in a core subunit of histone H3 lysine 4 methyltransferase complexes, which is required for chromatin-mediated silencing (Jeong et al., 2002; van Dijk et al., 2005). *Chlamydomonas* strains where DCL1 expression was suppressed by transgenic RNAi in a wild type background (CC-124) or in the Mut11 mutant background have also been described (Casas-Mollano et al., 2008). For the isolation of AGO1-associated sRNAs, we fused the FLAG tag (Einhauer and Jungbauer, 2001; Voshall et al., 2015) to the N-terminal end of the Chlamydomonas *AGO1* (*Cre02.g141050*) coding sequence. This construct was placed under the control of *PsaD* regulatory sequences and transformed into Maa7-IR44 by electroporation (Yamano et al., 2013). The *PsaD* regulatory sequences have been reported to allow reliable expression, with reduced incidence of gene silencing, of cDNA transgenes in *C. reinhardtii* (Fischer and Rochaix, 2001).

Unless noted otherwise, *C. reinhardtii* and other algal species were routinely grown in tris-acetate-phosphate (TAP) medium (Harris, 2009). For nitrogen deprivation analyses, cells initially grown in nutrient replete TAP medium to the middle of the logarithmic phase (OD_750_ ∼0.3-0.4) were collected by centrifugation, washed twice, and resuspended at a density of ∼1.0 x 10^6^ cells mL^-1^ in the same medium with or without (i.e., lacking ammonium chloride) nitrogen (Harris, 2009; Voshall et al., 2017). After 24 h of incubation under continuous illumination (∼150 μmol m^-2^ s^-1^ photosynthetic photon flux density), cells were harvested and immediately frozen in liquid nitrogen for subsequent RNA isolation. A similar protocol was used for the analysis of phosphate or sulfur deprived cells, following prior specifications (González-Ballester et al., 2010; Chávez Montes et al., 2014; Voshall et al., 2017). For heat-shock treatments, mid-log phase cells in TAP medium were incubated in a water bath at 42°C for 20 min.

### *AGO1* gene editing

Following a previously described protocol (Akella et al., 2021), cells of the wild type CC-124 strain were electroporated with an *in vitro* assembled CRISPR/Cas9 ribonucleoprotein, targeting the *AGO1* exon1, and a modified single-stranded oligodeoxynucleotide, to serve as template for homology directed DNA repair. After a 48-h recovery period, electroporated cells were spread on TAP agar plates containing oxyfluorfen (Akella et al., 2021). Surviving colonies were screened by PCR amplification (Cao et al., 2009) of the *AGO1* target site, using a primer (AGO1-mut-F1) that anneals exclusively to the edited sequence. The oligonucleotides and primers used for these experiments are listed in Supplemental Table S5.

### Isolation of AGO1-associated sRNAs, library preparation, and sequencing

FLAG-tagged AGO1 was affinity purified from cell lysates as previously described (van Dijk et al., 2005; Voshall et al., 2015). RNAs associated with AGO1 were purified with TRI reagent (Molecular Research Center) and contaminant DNA was removed by DNase I treatment (Ibrahim et al., 2010; Ma et al., 2013). Two independent small cDNA libraries were prepared with the small RNA v1.5 sample prep kit (Illumina), following the manufacturer’s protocol, and sequenced with a Hiseq1500 (Illumina). All sequencing data were deposited into the NCBI Sequence Read Archive (SRA) under BioProject PRJNA765360.

### sRNA mapping and profiling

Adaptor-trimmed reads were mapped, by using Bowtie2 (version 2.2.9) with default parameters (Langmead and Salzberg, 2012), to the unmasked *C. reinhardtii* genome v5.5 downloaded from Phytozome version 12.1 (https://phytozome.jgi.doe.gov/pz/portal.html) (Goodstein et al., 2012). Mapped reads were filtered to remove those shorter than 18 nt or showing alignments with gaps or mismatches. The population of reads that mapped perfectly to the genome was then profiled based on length. Genome-mapped reads were also classified as matching the chloroplast or the mitochondrial genomes, ribosomal RNAs (rRNAs), small nuclear or nucleolar RNAs (snRNAs or snoRNAs), transfer RNAs (tRNAs), or transposable elements. All sequences except for transposable elements were taken from Genbank (accession numbers BK000554.2 and NC_001638.1 for the chloroplast and mitochondrial genomes, respectively). Transposon sequences were taken from Repbase (Kapitonov and Jurka, 2008). After filtering the above sequences, the remaining mapped reads were further classified based on their match to structural categories in the annotated *Chlamydomonas* nuclear genome (Creinhardtii_281_v5.5.gene.gff3), such as intergenic or genic regions, the latter partitioned in 5’ UTRs, exons, introns, or 3’ UTRs. The expression level in reads per million (RPM) for each mapped sRNA was determined by the formula: RPM = [(10^6^*C*)/*N*], where *C* is the number of mapped reads corresponding to an individual sRNA sequence in the library and *N* is the total number of mapped reads in the library (Voshall et al., 2015). For comparative analyses, publicly available small RNA sequencing data from wild-type *C. reinhardtii* and a *dcl3* null mutant (Valli et al., 2016) were retrieved from the NCBI Sequence Read Archive (Project accession number PRJEB10672). AGO3-associated small RNA sequencing data (Voshall et al., 2015) were also retrieved from NCBI SRA (SRR1747077).

### Genome clustering of AGO1-associated sRNAs and secondary structure prediction of putative RNA precursors

After removing reads that mapped to known noncoding RNA categories, transposons, the chloroplast or the mitochondrial genomes, the remaining sRNA reads were clustered by genomic location such that there was no more than 100 nt between adjacent reads, regardless of strand, in a given cluster. The reads on both strands in the same genomic location were placed in the same cluster. Then the genomic sequence for each strand of a cluster was folded using version 2.3.1 of RNAfold from the Vienna RNA package (Lorenz et al., 2011), to assess whether putative precursor RNAs for the sequenced small RNAs may have secondary structures. Clusters were also manually examined to assess processing accuracy of the 5’ and 3’ ends of the predominant read(s), the frequency of the predominant read(s), and the extent of complementarity (i.e., dsRNA stem formation) between the two arms of predicted hairpins.

### Analysis of sRNA size distribution in green algae and land plants

Publicly available total small RNA sequencing data corresponding to whole cells or whole organisms (in non-reproductive phases) were retrieved from the NCBI Sequence Read Archive for the following species: *Volvox carteri* (SRR1029443), *Physcomitrium patens* (SRR1842137), *Marchantia polymorpha* (SRR2179617), *Picea abies* (SRR14056790), *Ginkgo biloba* (SRR1658901), *Zea mays* (SRR895785), and *Arabidopsis thaliana* (SRR4124967). The 3’ end adaptor sequences were trimmed using the fastx toolkit (http://hannonlab.cshl.edu/fastx_toolkit/), and the reads were then mapped to the respective reference genomes as described above. Genome-mapped redundant reads, longer than 17 nts, were then profiled based on their size.

### Prediction of *Chlamydomonas* long-sRNA targets

Putative long sRNA target transcripts were predicted using psRNATarget schemaV2 (2017 release) (Dai et al., 2018). Default parameters were used except for the expectation, which was set to 4 (instead of the default 5) to increase stringency. The transcript library used in the search was ‘cDNA library Phytozome 11, 281_v5.5.’ Putative functions of the predicted targets were evaluated by using the annotations of *Chlamydomonas* genes [if available through Phytozome v. 12.1, (Goodstein et al., 2012)] as well as conserved protein domains.

### Differential expression analysis of predicted long-sRNA targets under various nutritional conditions

Publicly available poly(A) mRNA sequencing data collected from wild type *C. reinhardtii* grown under different nutritional conditions were retrieved from the NCBI Sequence Read Archive for the following treatments: phosphorus deprivation (SRR1216592, SRR1216593, SRR1216594, SRR1216595), sulfur deprivation (SRR1521728, SRR1521729, SRR1521742, SRR1521743) and nitrogen deprivation (SRR1521680, SRR1521684, SRR1521722, SRR1521723). Standard RNA-seq analysis tools, Tophat and Cuffdiff, were used to compare gene expression under nutrient-replete or nutrient-deplete conditions (Trapnell et al., 2009). Transcript abundance was analyzed as Reads Per Kilobase of transcript per Million mapped reads (RPKM), which normalizes read counts based on both transcript length and total number of reads, using the formula: RPKM = [(10^9^*C*)/(*NL*)], where *C* is the number of reads mapped to each transcript, *N* is the total number of mapped reads in the library, and *L* is the transcript length in nucleotides (Mortazavi et al., 2008).

### Phylogenetic analysis

Argonaute protein sequences from Viridiplantae were obtained from the NCBI database and the Phycocosm algal portal (https://phycocosm.jgi.doe.gov/phycocosm/home). Polypeptides homologous to AGO-PIWI were identified by BLAST or PSI-BLAST searches of protein and/or translated genomic DNA sequences (see Supplemental Figure S1 for their accession numbers). The Phylogeny.fr server (Dereeper et al., 2008) was used to perform protein sequence alignment with MUSCLE (Edgar, 2004) and maximum-likelihood phylogeny reconstruction with PhyML (Guindon et al., 2010). Bootstrap analysis was performed with 500 pseudoreplicates. Visualization of the phylogeny was done using TreeDyn (Chevenet et al., 2006).

### Calculation of Ka/Ks ratios

Orthologous candidates for the *AGO1* and *AGO2/AGO3* genes (Supplemental Table S4) were identified from *C. reinhardtii* and closely related species, including *Chlamydomonas incerta*, *Chlamydomonas schloesseri, Edaphochlamys debaryana*, and *Chlamydomonas eustigma*, based on the phylogeny shown in Supplemental Figure S1. Multiple sequence alignment of each orthologous protein set was performed using the E-INS-i algorithm of MAFFT (Katoh et al., 2019). The nucleotide sequences coding for the AGO1 and AGO2/AGO3 proteins were aligned based on the protein alignments using TranslatorX (Abascal et al., 2010). The ratios of nonsynonymous (Ka) to synonymous (Ks) nucleotide substitution rates among these coding sequences were estimated by using the Ka/Ks Calculation tool available at the Norwegian Bioinformatics platform (http://services.cbu.uib.no/tools/kaks). It uses the methods found in Liberles (2001) and Siltberg and Liberles (2002). For each dataset, the coding sequence alignment produced using MAFFT was provided, and the maximum likelihood phylogenies (with the discrete Zhang substitution matrix option) were reconstructed. The boxplot of the results was generated using R (https://www.R-project.org/).

### RNA analyses

Total RNA was purified with TRI reagent (Molecular Research Center), in accordance with the manufacturer’s instructions (Ibrahim et al., 2010; Ma et al., 2013), from *C. reinhardtii*, *C. incerta*, or *E. debaryana* cells grown under the different trophic conditions. For sRNA northern blot analyses, total RNA samples were resolved in 15% polyacrylamide/7M urea gels and electroblotted to Hybond-XL membranes (GE Healthcare) (Ibrahim et al., 2010; Ma et al., 2013). Blots were hybridized with ^32^P-labeled oligo-DNA probes using the High Efficiency Hybridization System (Molecular Research Center) at 40°C for 48 h (Ibrahim et al., 2010; Ma et al., 2013). Specific sRNAs were detected by hybridization with DNA oligonucleotides labeled at their 5’ termini with [γ-^32^P]ATP and T4 Polynucleotide Kinase (New England Biolabs) as previously described (Ibrahim et al., 2010; Ma et al., 2013). The oligonucleotides used as hybridization probes are listed in Supplemental Table S5.

For reverse transcriptase (RT)-PCR analyses, reverse transcription reactions were performed as previously described (Carninci et al., 1998) using Superscript^TM^ III (Invitrogen). The synthesized cDNA was then used as a template in standard PCRs (Sambrook and Russell, 2001) using a number of cycles showing a linear relationship between input RNA and the final product, as determined in preliminary experiments. The PCR conditions for amplification of the Actin control were 25 cycles at 94 °C for 30 s, at 58 °C for 30 s and at 71 °C for 30 s. The PCR conditions for amplification of the *DCL2* or *Cre03.g164550* (*BOLA-LIKE*) genes were 26 to 30 cycles at 94 °C for 30 s, at 60 °C for 30 s and at 71 °C for 30 s. Aliquots (8 µL) of each RT-PCR were resolved on 1.5% agarose gels and visualized by ethidium bromide staining (Sambrook and Russell, 2001). All primers used for RT-PCR are listed in Supplemental Table S5.

## Accession numbers

All sequencing data generated in this study have been deposited in the National Center for Biotechnology Information Sequence Read Archive under BioProject PRJNA765360. Accession numbers of public datasets analyzed in this work are listed in Materials and methods under the appropriate sections.

## Supplemental data

The following materials are available on the online version of this article.

**Supplemental Figure S1.** Maximum-likelihood phylogeny among AGO homologs from Viridiplantae.

**Supplemental Figure S2.** Abundance of long sRNAs in wild type and *ago1* mutant strains of *Chlamydomonas reinhardtii*.

**Supplemental Figure S3.** Secondary structure plots of predicted precursor RNAs for clusters of AGO1-associated long sRNAs.

**Supplemental Figure S4.** Comparison of the structure of *FRA2/BolA-like protein 2* gene orthologs in algae of the Trebouxiophyceae and Chlorophyceae classes.

**Supplemental Figure S5.** VISTA plot of nucleotide conservation of the *Chlamydomonas incerta FRA2/BolA-like protein 2* gene (g5696.t1) relative to its *C. reinhardtii* ortholog.

**Supplemental Figure S6.** VISTA plot of nucleotide conservation of the *Chlamydomonas schloesseri FRA2/BolA-like protein 2* gene (g16856.t1) relative to its *C. reinhardtii* ortholog.

**Supplemental Figure S7.** Long sRNAs in Chlamydomonas reinhardtii, Chlamydomonas incerta, and Edaphochlamys debaryana.

**Supplemental Figure S8.** DCL2 transcript and protein abundance in *Chlamydomonas reinhardtii*.

**Supplemental Figure S9.** Ratios of nonsynonymous (Ka) to synonymous (Ks) substitution rates calculated from Actin, AGO1, and AGO2/AGO3 coding sequences among *C. reinhardtii* and four closely related algal species.

**Supplemental Figure S10.** Binding site of Cluster 51304+ long sRNAs on the 3’ UTR of the *Cre03.g164550* (*FRA2/BolA-like protein 2)* transcript.

**Supplemental Table S1.** Abundance of AGO1-associated long sRNAs and miRNAs in *Chlamydomonas reinhardtii* cultured under nutrient replete or nitrogen deprived conditions.

**Supplemental Table S2.** Predicted targets of the most abundant AGO1-associated long sRNAs from cluster 51304+.

**Supplemental Table S3.** Differential expression analysis of predicted target transcripts of cluster 51304+ long sRNAs under sulfur, nitrogen or phosphorus deprivation.

**Supplemental Table S4.** List of accession numbers for the coding sequences used for the Ka/Ks calculation.

**Supplemental Table S5.** Oligonucleotides and primers used in this study.

## Acknowledgements

We are thankful to members of the Cerutti and Moriyama lab for helpful technical suggestions on sRNA northern blotting.

## Authors contributions

E.M. and H.C. conceived and designed the project. Y.L., E.J.K., and A.V. performed the experiments. Y.L., A.V., E.M., and H.C. implemented software and analyzed the data. Y.L., E.J.K., E.M., and H.C. wrote and edited the manuscript. All authors read and approved the final article.

